# Where does time go when you blink?

**DOI:** 10.1101/506279

**Authors:** Shany Grossman, Chen Guata, Slav Pesin, Rafael Malach, Ayelet N. Landau

## Abstract

Retinal input is frequently lost due to eye blinks, yet humans rarely notice these gaps in visual input. While previous studies focused on the psychophysical and neural correlates of diminished awareness to blinks, the impact of blinks on the perceived time of concurrent events is unknown. Here, we investigated whether the subjective sense of time is altered by spontaneous eye blinks, and how this link may inform mechanisms of time perception. We found that participants significantly underestimated the duration of a visual stimulus when a blink occurred during the stimulus. Importantly, this effect was not present when durations of an auditory stimulus were judged. These results point to a link between spontaneous blinks, previously demonstrated to induce suppression of activity in early visual cortex, and a compression of subjective time. The findings suggest that ongoing encoding within modality-specific sensory cortices, independent of conscious awareness, inform the subjective sense of time.

---

Spontaneous eye blinks trigger an occlusion of retinal input for a considerable duration, from tens to hundreds of milliseconds (VanderWerf, Brassinga, Reits, Aramideh, & Ongerboer de Visser, 2003). Nevertheless, these frequent interruptions usually go unnoticed as our visual experience remains continuous. Diminished awareness to visual stimulation during blinks was previously validated in a psychophysical study that found a significant reduction in visual sensitivity during voluntary eye blinks (Volkmann, Riggs, & Moore, 1980). This suppression of visual detection thresholds during blinks occurred when light was delivered through the roof of the mouth and was therefore independent of eyelid position. Subsequent electrophysiological studies in cats (Buisseret & Maffei, 1983) and in primates (Gawne & Martin, 2000, 2002) reported a decrease in firing rates of neurons in the primary visual cortex during blinks. These studies also supported a neural, rather than an optical, source of blink-related reduction in visual activity. Human studies have provided further support of an extra-retinal suppression in early visual regions during voluntary blinks, using fMRI (Bristow, Haynes, Sylvester, Frith, & Rees, 2005), as well as suppression of transient activity in the visual cortex during spontaneous and voluntary blinks, as revealed in intracranial EEG recordings (Golan et al., 2016).

Given the link between blinks and suppression of both visual sensitivity and neural activity, a natural yet unexplored question concerns what happens to subjective time during spontaneous eye blinks. This question offers a unique opportunity to study the link between mechanisms of time perception and continuous processing of sensory input. The neural underpinnings of time perception are an active, unresolved field (Ivry & Schlerf, 2008; Wittmann, 2013). *Dedicated models* for timing postulate that duration estimation is implemented by a neural mechanism that is fully designated to timing, for example, by postulating a neural pacemaker. In contrast, *intrinsic models* assume that time is inherently encoded by the neural resources invested in sensory processing. Whether neural mechanisms of timing are modality specific is somewhat related to the distinction between dedicated and intrinsic models of timing: intrinsic models would postulate timing as a modality specific computation. Investigating the link between suppressed neural processing – indexed by spontaneous eye blinks – and sensory duration judgments can provide important evidence for the role of ongoing perceptual encoding in informing temporal estimation.

Here we investigated how spontaneous eye blinks impact the subjective sense of time. An eye blink during a visual stimulus leads to an unnoticed gap in retinal input. Importantly, we used both visual and auditory stimuli as timed intervals in two separate experiments in order to test whether time perception was affected by the input loss in a modality specific manner.

## Methods and Materials

### Participants

A total of 29 and 30 participants took part in the visual and the in the auditory experiments, respectively. Inclusion criterion was defined prior to data analysis as having a minimum of 10 percent blink and blink-free intervals. In both the visual and the auditory experiments, 7 individuals did not meet this inclusion criterion and were therefore discarded from all analyses. The analyses reported in this paper were thus carried out on 22 visual participants (14 females, 24±2 years old) and 23 auditory participants (17 females, 22.5±3 years old). Only for the individual bisection point analysis (**Fig. 2c**) did we apply further exclusion criteria whereby participants with a poor fit were discarded (see **Psychometric function fit**). A total of 18 and 15 participants in the visual and auditory experiments, respectively, (4 and 7 participants excluded, respectively) were therefore included in this analysis. Written informed consent was obtained from all participants in line with the institutional IRB approval from the Hebrew University of Jerusalem.

**Figure 1.**
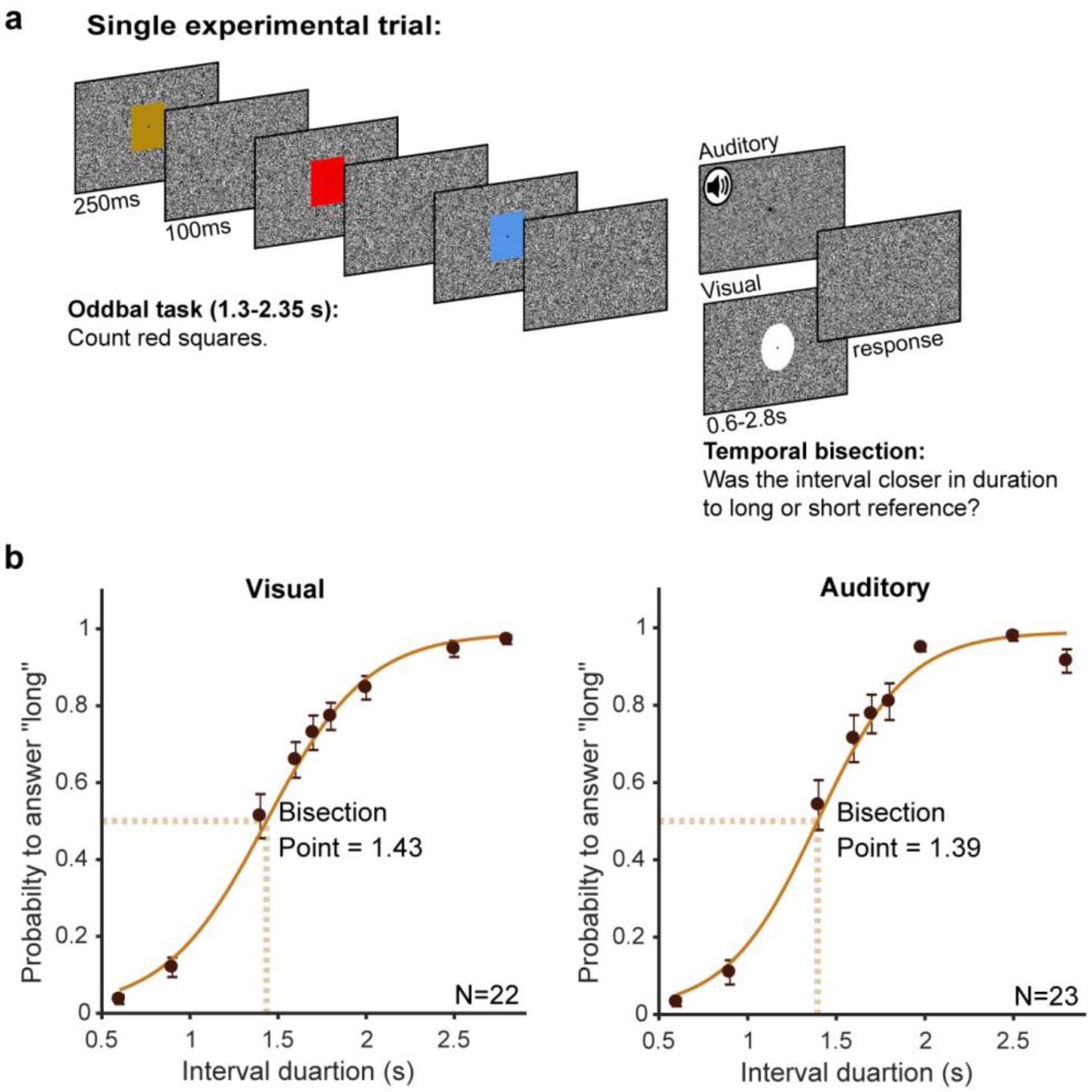
Experimental design and group performance for both blink and blink-free intervals. **a)** Schematic illustration of a single trial. Each trial consisted of two combined sub-tasks appearing in continuous succession. The oddball task was emphasized as the main task, whilst temporal bisection was defined as secondary. A white central disc and a segment of white noise were used as the timed interval in the visual and auditory experiments, respectively. On each temporal bisection trial, the timed stimulus was presented for one of 9 predefined durations (ranging from 0.6 to 2.8 sec). Participants judged whether the temporal interval was closer to the short or long reference interval, with which they were familiarized during the initial stage. During the oddball task, central colored squares were flashed for 250 ms with a fixed ITI of 100 ms. Participants were instructed to count the number of red squares that appeared on every experimental block (4 blocks in total, each consisting of 100 trials). **b)** Group psychophysical performance derived from all intervals (blink and blink-free) of all participants. Group psychometric functions and bisection points (BP) for each modality were estimated by a logistic fit to the mean probabilities to answer long across subjects. They are presented here for visualization only, note that all statistical tests took a within subject approach. Error bars denote ±1 SEM.

**Figure 2.**
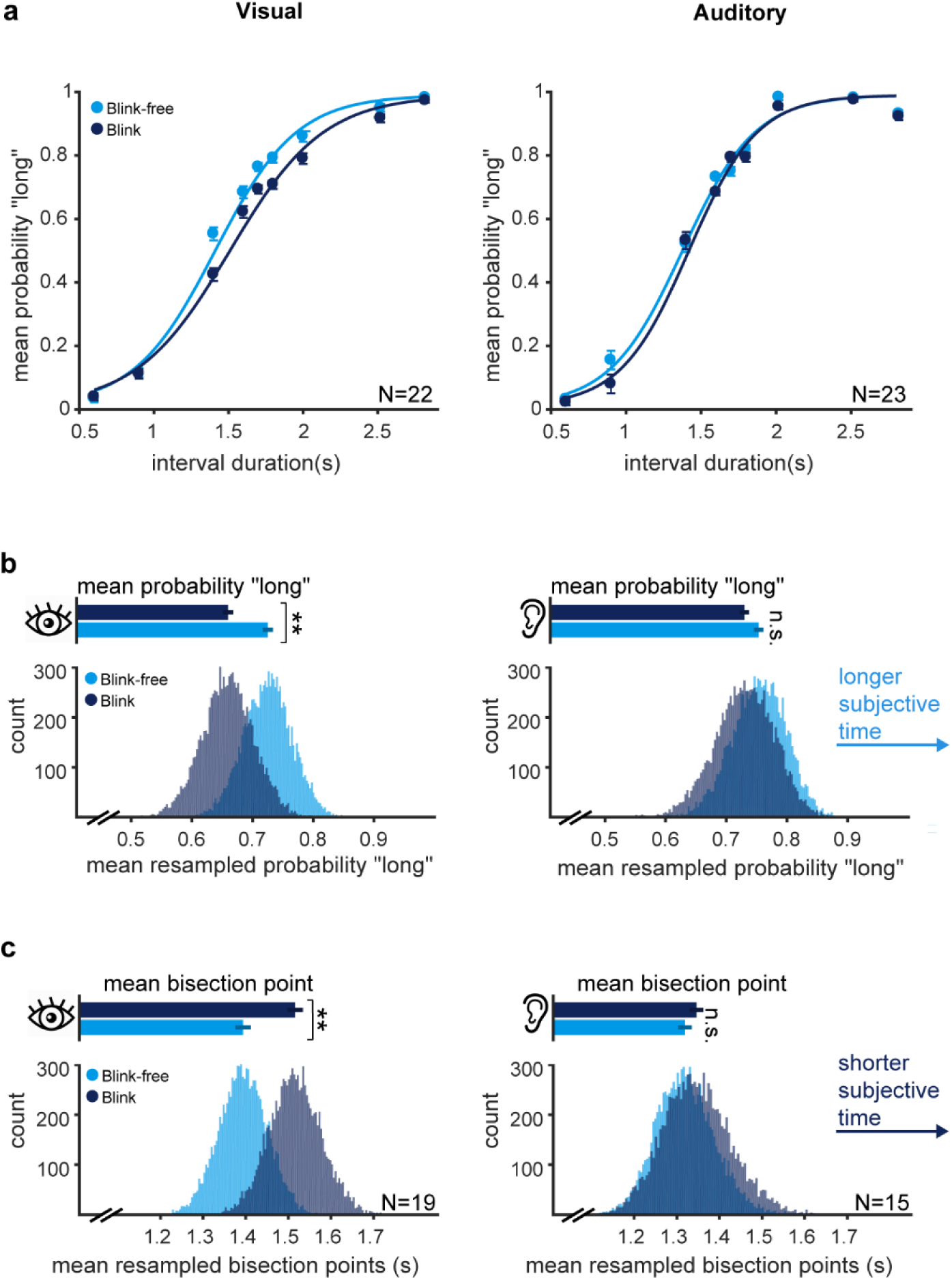
Temporal bisection performance in blink and blink-free intervals. **a)** For each of the 9 possible interval durations (0.6 0.9 1.4 1.6 1.7 1.8 2 2.5 2.8), the mean probability to judge the duration as closer to the long reference interval is presented separately for blink and blink-free intervals. Results are presented separately for visual (left panel) and auditory (right panel) timed intervals. Note the visual-specific modulation of performance in blink intervals relative to the blink-free intervals. **b)** Histograms describe bootstrap distributions of the mean probability to judge a visual (left panel) or an auditory (right panel) interval from the dynamic range (5 intermediate levels) as long in the presence and in the absence of blinks. Top horizontal bars denote the original mean probabilities for blink and blink-free intervals. Note that higher values for mean probability “long” reflect longer subjective time. **c)** Individual BPs are larger in the presence of spontaneous blinks. Histograms describe bootstrap distributions of mean bisection point as derived from individual psychometric function for blink and blink-free intervals, separately. Note that larger BP corresponds to shorter subjective time. All error bars in the figure denote ±1 SEM following Cousineau-Morey correction for a within participant design(Morey, 2008). Figure design inspired by (Terhune, Sullivan, & Simola, 2016).

### Stimuli

The experiment was programmed in MATLAB (2017a, MathWorks) using Psychtoolbox-3(Kleiner et al., 2007). Oddball stimuli were 10 colored squares. For the visual experiment, a white central disc was displayed for marking the timed interval. Both the squares and the disc subtended a visual angle of 11.4°. For the auditory experiment, bursts of white noise generated in MATLAB were delivered through headphones for marking the timed intervals. Throughout both visual and auditory experiments, a random noise black and white pattern was displayed as a background and a central red fixation point was presented.

### Experimental Procedures

The experimental session began with a familiarization stage aimed to introduce the participant with the duration of the short (0.6 sec) and long (2.8 sec) reference intervals. The reference intervals were presented and the participant could replay them as many times as he/she felt needed.

The main experiment was a dual task: each trial (400 trials in total) consisted of an oddball part and a temporal bisection part (see **Fig. 1a** for a schematic illustration). A trial began with a jittered sequence of 4-7 colored central squares, flashed for 250 ms each with a fixed ITI of 100 ms. The participant was instructed to report how many red squares appeared during the entire oddball sequences of the experiment. Immediately upon termination of the oddball part (i.e. without any time delay), the timed interval was presented for one out of 9 possible durations spanning the range between the two extreme reference intervals: 0.6 0.9 1.4 1.6 1.7 1.8 2 2.5 2.8 (sec). Upon the offset of the timed interval, participants were instructed to indicate by pressing one of two possible keys whether the current interval was closer in its duration to the short, or long, reference interval. Response time was unlimited and response collection initiated the next oddball sequence (i.e. the beginning of a new trial). All participants completed a short training prior to the main experiment. To maximize the probability that spontaneous blinks would occur during timed intervals, oddball task was emphasized as the main task whilst bisection was defined as secondary.

### Equipment

We used a video-based eye tracker (Eyelink 1000, SR Research) to monitor continuously eye position and pupil diameter of participants. Eye data was recorded binocularly at a sampling rate of 1000 Hz. Messages were sent from the experiment PC to the Eyelink PC upon stimuli presentation for offline alignment of the experimental log to eye tracking time series.

### Classification of blink and blink-free intervals

The raw pupil time series of each participant was processed offline in a semi-automatic blink detection procedure, composed of two stages. First, blinks were automatically marked as all segments with missing pupil samples, with blink onset corresponding to the point of maximal acceleration in the decrease of pupil size, and blink offset corresponding to maximal de-acceleration point in increase of pupil size. These points should reflect the time point in which the eyelid started covering the pupil and the time point in which the pupil was fully exposed again, respectively. Second, we manually inspected the output of the automatic detection on top of the raw pupil-size time series and disqualified segments without a gradual decrease and increase in pupil size surrounding the putative blink. Disqualified segments, as well as segments that exceeded 400 ms, were logged and excluded from all analyses.

Following temporal alignment of pupil time series to the experimental log, we divided timed intervals into blink and blink-free. Importantly, blink intervals were defined only when a blink occurred entirely within the interval - its onset detected at least 50 ms post the interval’s onset, and its offset preceding the interval’s offset in at least 50 ms. Timed intervals in which a blink overlapped with their onset or offset were logged and excluded from all analyses.

### Psychometric function fit

Individual psychometric functions were estimated through a logistic function fit based on a maximum likelihood criterion and executed by scripts available in the Palamedes toolbox (Prins, 2014). Two free parameters were estimated - the threshold (i.e. bisection point) and the slope, while the guess and lapse rate were fixed and predefined as equal to 1% each. A measure of goodness of fit, pDev, was estimated for each fit. To estimate pDev, deviance values of simulated data drawn from the best-fitting psychometric curve were computed over 1000 iterations. The proportion of simulated deviance values that were greater than the deviance value of the original data corresponds to the pDev value. Thus, larger values of pDev reflect a better fit to the measured data, while pDev < 0.05 indicates a poor fit of the data(Kingdom & Prins, 2010). Since the individual bisection point (BP) analysis (**Fig.1c**) relied on psychometric fits, participants with poorly fitted functions (pDev<0.5) were excluded from this analysis. Two additional participants, one from the auditory and one from the visual experiment, had no blink intervals in the two shortest levels (0.6 and 0.9 sec) and were therefore excluded from individual fitting analysis as well. Thus a total of 3 and 7 participants were excluded from the visual and auditory experiments, respectively, for the individual BP analysis. Notably, the results of the this analysis remained unchanged also upon inclusion of poor fitted participants (paired t-test, visual: p=0.007, t(20)=3.02; auditory: p=0.54, t(21)=0.62).

Group psychometric curves presented in **Fig. 2a** and **Fig. 2b** were estimated by fitting a logistic function to the group mean proportion “long” in each interval duration. Importantly, this fit is purely for visualization and does not contribute to any of the analyses presented here. The main analyses presented in the results section, whether on probability “long” values or on BPs, took a within subject approach with subjects as a random effect.

### Control analysis for unequal samples size

Naturally, the number of blink- and blink-free intervals was not matched within a participant or a specific interval level, with a bias towards having fewer blink intervals. We therefore ran a control analysis to ensure this bias did not affect the observed results. To this aim, we randomly subsampled the data such that each interval duration of each participant had an equal amount of blink and blink-free intervals. This essentially means taking a random subsample of the intervals in the condition (blink/blink-free) with the larger n. We then repeated the analysis presented in **Fig.1a**, performing a paired t-test on the proportion “long” across the dynamic range in blink vs. blink-free intervals for this size-matched subsample of the data. On each iteration (n=1000), the analysis was carried out on a size-matched random subsample of the data, equating the number of blink and blink-free intervals for every participant and interval duration. The resultant p-value distributions are presented in **Fig. S1**. For the visual intervals, all resultant p-values were smaller than 0.05 (95th percentile = 0.004). In contrast, size-matched sub-samples of auditory intervals resulted in a wide range of p-values, 98% of them larger than 0.05 (95th percentile = 0.9).

## Results

A major challenge in studying the impact of spontaneous blinks on duration estimation is that a single-task approach leads to very few blinks during timed intervals, as participants naturally tend to blink between the to-be timed intervals. We therefore combined a temporal bisection task in a dual task paradigm. Upon mastering discrimination between long and short reference intervals (0.6 and 2.8 sec, respectively), participants proceeded to the main experiment. Each trial consisted of two successive sub tasks: an oddball detection task during a rapid serial visual presentation (RSVP) followed by a temporal bisection task (see **Fig. 1a** for a scheme of the experimental design). On each temporal bisection trial, a timed interval was displayed for one out of nine possible durations, ranging from 0.6 to 2.8 sec. Participants were instructed to report whether they perceived the interval as being more similar to the long, or short, reference interval. Temporal bisection was performed on visual or auditory intervals in two separate experiments, with all timing parameters held constant. Visual-timed intervals consisted of a central white disc subtending 11.4° visual angle, whereas auditory-timed stimuli were intervals of white noise.

Prior to inspecting the impact of blinks on duration estimation, we first observed the overall performance of the group by collapsing data from all participants and plotting proportion “long” responses as a function of stimulus duration (**Fig. 1b**). Based on individual logistic fits to all intervals (blink and blink-free) of each participant, we estimated the mean bisection point (BP) across participants. Interestingly, the mean BP was significantly shorter than the objective mid-point between the two extreme reference intervals (1.7 sec) and was estimated to be 1.4 sec (95% CI: 1.28-1.51) and 1.39 sec (95% CI: 1.27-1.5) for visual and auditory intervals, respectively. This bias is in line with a previous meta-analysis linking intervals spread greater than two (quantified as the long reference/short reference ratio) with an underestimation of the true bisection point (Kopec & Brody, 2010) (see reference for a suggested model which accounts for this bias).

For the main analysis, timed intervals were classified as either blink or blink-free. Blink intervals were defined as consisting of a full blink, which was initiated at least 50 ms after the onset of the interval and terminated at least 50 ms prior to its offset. Twenty-two and 23 subjects who participated in the visual and auditory experiments, respectively, met our predefined inclusion criteria and were included in the analyses (see Methods for inclusion criteria). On average, participants from the visual and auditory experiments had 126.7±69.3 blink trials (38.3±24% of the total trial count). There was no significant difference in the percent of blink intervals between the visual and auditory experimental groups (independent samples t-test, t(43)=1.009, p=0.32).

In order to examine the impact of spontaneous blinks on duration judgments, we took two complementary approaches: a direct quantification of individual performance and an estimation of individual bisection points (BP) through a psychometric function fit. Accordingly, we first computed the probability to answer “long” at each interval duration and condition (blink vs. blink-free) per participant. As presented in **Fig. 2a**, this analysis revealed a clear reduction of the probability to judge visual intervals as “long” in the presence of a spontaneous blink, as compared with the same probabilities for blink-free trials. The probability to answer “long” across the dynamic range (collapsing 5 intermediate levels for which judgment is more difficult, 1.4-2 sec) was significantly smaller in blink, as compared to blink-free, intervals (**Fig. 2b**; paired t-test, t(21)=3.8, p=0.001, Cohen’s d = 0.81). Here too, the effect was specific to the visual modality as performance did not differ between blink and blink-free auditory intervals (paired t-test, t(22)=1.52, p=0.14).

We next compared the bisection point (BP) between blink and blink-free data. The BP corresponds to the interval duration which is equally likely to be judged as “long”, or as “short”. To this end, we fit a psychometric function to each individual’s performance, for blink and blink-free intervals separately, and derived the BP from the estimated function (see Methods). For visual intervals, we found a significant increase in BPs for intervals containing blinks as compared to blink-free intervals (**Fig. 2c;** paired t-test: t(18)=3.4, p=0.003, Cohen’s d=0.77), further indicating a time compression with blinks. It is noteworthy that the group average shift in BP (121±36 SEM) was similar in magnitude to the average blink duration (112±9 SEM). Here too the effect was evident only for visual intervals, as we did not observe any difference in BPs in the auditory experiment (paired t-test: t(14)=1.29, p=0.22). Overall, there were significantly less blink intervals than blink-free intervals in both the visual and the auditory groups (paired t test; visual: t(21)=3.36, p=0.003; auditory: t(22)=2.22, p=0.036). In order to rule out the possibility that differences in trial-count had contributed to the observed underestimation for blink-intervals, we repeated the main analyses, presented in **Fig. 2a-b,** using a stratification approach that consisted of iteratively subsampling the data to achieve an equal number of blink and blink free interval per participant and duration (see Methods for details). The results of this analysis rule out smaller trial-count of blink intervals as a confounding factor.

Since spontaneous blinks may also be coupled with attentional lapses, a possible explanation of the observed time compression may be attenuated attention. The specificity of the effect to the visual modality goes against such an attentional account. Nonetheless, to further test this possibility, we compared the reaction times between blink and blink-free intervals. In both the visual and auditory data, reaction times were not significantly longer following blink intervals, as compared to blink-free intervals (paired t-test, visual: t(21)=0.72; p=0.48; auditory: t(22)=1.51, p=0.15), arguing against decreased attention in blink intervals in either one of the modalities.

Finally, it could be argued that blinks during auditory intervals did not perturb duration judgments since audition outperforms vision with regards to subsecond temporal resolution (Ortega, Guzman-Martinez, Grabowecky, & Suzuki, 2014). However, we found no difference between individual bisection points estimated on blink-free auditory intervals and blink-free visual intervals (independent samples t-test, t(36)=0.89, p=0.38), or between the corresponding slopes of the psychometric functions (independent samples t-test, t(36)=-1.3, p=0.19). This counters the possibility that differences in task difficulty or performance precision underlie the modality specificity aspect of the current finding.

## Discussion

The current results reveal that duration estimation of visual, but not auditory, input is significantly reduced when a blink occurs within the estimated time interval. A converging line of studies has shown that blinks induce a momentary reduction in visual sensitivity and a suppression of transient neural activity in the visual cortex, both originating in an extra-retinal signal. The current findings demonstrate that the unaware loss of visual sensitivity due to blinks is coupled with a loss of subjective time of an equivalent duration. Importantly, the specificity of the effect to the visual modality supports the view whereby the amount of neural processing invested in sensory encoding of an incoming stimulus is an integral part of temporal estimation. Thus, a reduction in visual processing, indexed by a blink, leads to the concurrent loss of subjective time.

Distortions of visual temporal estimation, at a shorter time scale than studied here, have been linked to another type of eye movement - large saccades. Findings pointing to pre-saccadic time compression (Morrone, Ross, & Burr, 2005; Terao, Watanabe, Yagi, & Nishida, 2008) and to post saccadic time dilation (Yarrow, Haggard, Heal, Brown, & Rothwell, 2001) have been proposed to relate to predictive shifts in spatial receptive fields. Specifically, underestimation of pre saccadic intervals has also been suggested to relate to precision errors in encoding the onset and offset of a stimulus (Terao et al., 2008), yet the impact of ongoing, cumulative, processing of the stimulus on time distortion has not been tested. Importantly, saccades, different from eye-blinks, contain a spatial element and are events tightly linked to visual awareness of the change in retinal input. Here the use of blinks allowed the direct examination of input loss in the absence of retinal motion, predicted or actual image displacement, awareness or any perceived discontinuity. Therefore, our data speak to a purely temporal consequence of eye blinks and the accompanying loss of sensory input at the supra-second time scale. We found that unconscious loss of visual input affects timing of visual intervals, but not auditory intervals, supporting a central role for ongoing sensory encoding in the subjective sense of time.

## Supplemental figures

**Figure S1.**
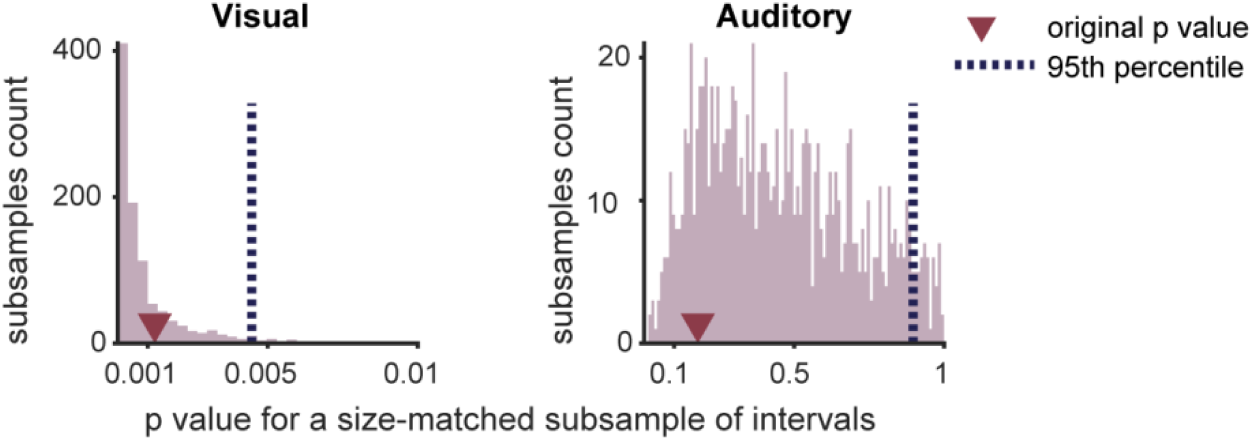
Control analysis for unequal number of trials between condition types. In order the ensure the effect was not driven by a smaller amount of blink intervals relative to blinks-free intervals, we repeated the analysis presented in **Fig.1b** 1000 times, each time subsampling the data to achieve a matched number of blink and blink-free interval in each individual and interval level. The histograms present the distribution of the resultant p values in the visual (right panel) and auditory (left panel) experiments.

## References

Bristow, D., Haynes, J.-D., Sylvester, R., Frith, C. D., & Rees, G. (2005). Blinking suppresses the neural response to unchanging retinal stimulation. Current Biology, 15(14), 1296–1300.

Buisseret, P., & Maffei, L. (1983). Suppression of visual cortical activity following tactile periorbital stimulation; its role during eye blinks. Experimental Brain Research, 51(3), 463–466.

Gawne, T. J., & Martin, J. M. (2000). Activity of primate V1 cortical neurons during blinks. Journal of Neurophysiology, 84(5), 2691–2694.

Gawne, T. J., & Martin, J. M. (2002). Responses of primate visual cortical neurons to stimuli presented by flash, saccade, blink, and external darkening. Journal of neurophysiology, 88(5), 2178–2186.

Golan, T., Davidesco, I., Meshulam, M., Groppe, D. M., Mégevand, P., Yeagle, E. M., et al. (2016). Human intracranial recordings link suppressed transients rather than ’filling-in’ to perceptual continuity across blinks. eLife, 5, e17243.

Ivry, R. B., & Schlerf, J. E. (2008). Dedicated and intrinsic models of time perception. Trends in cognitive sciences, 12(7), 273–280.

Kingdom, F., & Prins, N. (2010). Psychophysics: a practical introduction: Academic Press London.

Kleiner, M., Brainard, D., Pelli, D., Ingling, A., Murray, R., & Broussard, C. (2007). What’s new in Psychtoolbox-3. Perception, 36(14), 1.

Kopec, C. D., & Brody, C. D. (2010). Human performance on the temporal bisection task. Brain and cognition, 74(3), 262–272.

Morey, R. D. (2008). Confidence intervals from normalized data: A correction to Cousineau (2005). reason, 4(2), 61–64.

Morrone, M. C., Ross, J., & Burr, D. (2005). Saccadic eye movements cause compression of time as well as space. Nature neuroscience, 8(7), 950.

Ortega, L., Guzman-Martinez, E., Grabowecky, M., & Suzuki, S. (2014). Audition dominates vision in duration perception irrespective of salience, attention, and temporal discriminability. Attention, Perception, & Psychophysics, 76(5), 1485–1502.

Prins, N. (2014). Kingdom, FAA (2009). Palamedes: Matlab routines for analyzing psychophysical data.

Terao, M., Watanabe, J., Yagi, A., & Nishida, S. y. (2008). Reduction of stimulus visibility compresses apparent time intervals. Nature neuroscience, 11(5), 541.

Terhune, D. B., Sullivan, J. G., & Simola, J. M. (2016). Time dilates after spontaneous blinking. Current Biology, 26(11), R459–R460.

VanderWerf, F., Brassinga, P., Reits, D., Aramideh, M., & Ongerboer de Visser, B. (2003). Eyelid movements: behavioral studies of blinking in humans under different stimulus conditions. Journal of neurophysiology, 89(5), 2784–2796.

Volkmann, F. C., Riggs, L. A., & Moore, R. K. (1980). Eyeblinks and visual suppression. Science, 207(4433), 900–902.

Wittmann, M. (2013). The inner sense of time: how the brain creates a representation of duration. Nature Reviews Neuroscience, 14(3), 217–223.

Yarrow, K., Haggard, P., Heal, R., Brown, P., & Rothwell, J. C. (2001). Illusory perceptions of space and time preserve cross-saccadic perceptual continuity. Nature, 414(6861), 302.

